# DO ORGANIC, CONVENTIONAL, AND INTENSIVE APPROACHES IN LIVESTOCK FARMING HAVE AN IMPACT ON THE CIRCULATION OF INFECTIOUS AGENTS? A SYSTEMATIC REVIEW, FOCUSED ON DAIRY CATTLE

**DOI:** 10.1101/2024.02.20.581183

**Authors:** Massimo Pajoro, Matteo Brilli, Giulia Pezzali, Laura Kramer, Paolo Moroni, Claudio Bandi

## Abstract

A common thought is that extensive and organic breeding systems are associated with lower prevalence of infections in livestock animals, compared to intensive ones. In addition, organic systems limit the use of antimicrobial drugs, which may lead to lower emergence of antimicrobial resistances (AMR). To examine these issues, avoiding any a priori bias, we carried out a systematic literature search on dairy cattle breeding. Search was targeted to publications that compared different types of livestock farming (intensive, extensive, conventional, organic) in terms of the circulation of infectious diseases and AMR. A total of 101 papers were finally selected. These papers did not show any trend in the circulation of the infections in the four types of breeding systems. However, AMR was more prevalent on conventional dairy farms compared to organic ones. The prevalence of specific pathogens and types of resistances were frequently associated with specific risk factors that were not strictly related to the type of farming system. In conclusion, we did not find any evidence suggesting that extensive and organic dairy farming bears any advantage over the intensive and conventional ones, in terms of the circulation of infectious agents.

## 1. INTRODUCTION

It is a common view that intensive livestock farming facilitates the circulation of infectious agents within herds, possibly facilitating spill-over events of infections from animals to humans and the diffusion of emerging infectious diseases (EID) ^1^. Factors that are expected to facilitate the circulation of infections in intensive management are: high density and numerosity of animals; movement of animals (or animal-derived products) to food-processing industries (or markets); sub-optimal animal welfare conditions ^2^. However, this view has been called into question, at least for some types of farming management systems (FMS) ^1,3^. In addition, reliable opinions on this topic requires that intensive and extensive FMS are precisely defined ^1^. Intensive animal farming typically ensures a higher production per unit area ^1^. Intensive farming is thus expected to reduce the extension of the land used to support animal breeding, for a given amount of produce (e.g., milk, meat, eggs). Furthermore, intensive livestock farming is frequently based on large herds, with animals placed in dense aggregations, as opposed to extensive farming in which herds are generally small, and animals not over crowded ^4^. Therefore, according to ^1^, but differently from other views (e.g. ^4^), extensive farming increases the risk of circulation of infectious agents, since it is associated with the fragmentation of farming into small holdings, with higher number of animals in a given area, in order to obtain the same production that is achieved in intensive farming with fewer animals. The fragmentation of the farming system may increase contact with wildlife and the sylvatic environment. Thus, while it has been suggested that intensive farming is associated with increased circulation of infectious agents, an alternative hypothesis is that extensive breeding is more likely to favour infections in livestock. This issue can also regard organic farming, defined by specific requirements that differ among countries, and which shares some features with extensive systems, such as the space available to animals or the fact that high production yield is generally not a primary goal in either system. In addition, extensive farming is generally assumed to be associated with a high level of animal welfare, similar to organic farming ^5,6^. In turn, animal welfare is thought to be associated with increased resistance to infectious agents ^7^.

Along with the scientific literature, non-specialist publications (e.g. news publications, magazines and web sites devoted to scientific dissemination) have also discussed the “pros” and “cons” of intensive and conventional farming, at times in the absence of solid scientific evidence. The issue is thus well suited to be addressed through systematic analyses of the literature. A systematic review implies that search keys and inclusion and exclusion criteria are defined *a-priori*, in order to obtain unbiased retrieval of relevant publications. Systematic reviews are frequently performed according to the PRISMA protocol ^8^. The resulting scientific literature is then examined to build a general, unbiased view of the topic, rather than an “advocacy publication”.

In this systematic review we focused on dairy cattle farming, to evaluate the current knowledge on the circulation of infectious agents and the diffusion of antimicrobial resistance in relation to the type of management system. We specifically defined the search keys to retrieve publications in which the different types of managements (intensive and extensive; conventional and organic) have been directly compared. Furthermore, the circulation of infectious agents in herds, in terms of both acute and chronic infections, may impact on the quality of animal-derived foods. Therefore, a systematic review, or metanalysis, to determine whether the prevalence of infections in herd is influenced by the type of management systems might also provide indirect information on the quality of food production, and in this specific case, quality of milk and milk-derived products, as recently emphasized by the Food and Agriculture Organization of the United Nations^9^. The present metanalysis is not specifically addressed to livestock veterinary practitioners and stakeholders of milk production chain, whose interest is likely focused on specific risk factors associated with cattle infections. Rather, the aim is to target those “non-specialist” readers such as journalists, science communicators, and politicians, who have the responsibility to inform the general public, orienting (and taking) decisions on laws and regulations. Indeed, the issue of intensive and extensive production (or conventional and organic) is hotly debated in governmental institutions (e.g. ^10^), non-governmental organizations (e.g. ^11^), and newspapers (e.g. ^12^). In this context, the public deserves information based on unbiased reports to prevent the creation and circulation of ideological positions. The purpose of this article is thus to respond to this need for information, with a search on the scientific literature that is not biased by preconceptions.

## 2. METHODS

In this study, we followed the PRISMA guidelines ^8^ and carried out the bibliographic search on the Scopus database (search day: 4/3/2023). Literature retrieval strategy consisted in the search for publications in which dairy cattle management systems have been compared. Specifically, the type of searched comparisons was as follows: intensive VS extensive (IvsE) or conventional VS organic (CvsO). The different types of management systems were compared in relation to the following: 1) infectious agents (IA) or infectious diseases (ID); 2) antimicrobial resistance/susceptibility (AMR/AMS). The search terms were selected according to synonyms present in the literature. In addition, we assumed that publications focused on anti-parasitic and antimicrobial drug use (AMU) might provide results relevant to the issues of IA/ID and AMR/AMS, and we thus included terms related with the use of these drugs (even though the specific issue of AMU was out of the scope of this study). The following search terms and Boolean operators were used; the two query strings are reported below. Two query strings were used rather than combining all search terms into a single string in order to obtain two separate outputs for IvsE and CvsO.

( ( TITLE-ABS-KEY ( intensive* ) AND TITLE-ABS-KEY ( cow ) OR TITLE-ABS-KEY ( cattle ) AND TITLE-ABS-KEY ( milk ) OR TITLE-ABS-KEY ( dairy ) OR TITLE-ABS-KEY ( cheese ) AND TITLE-ABS-KEY ( infect* ) OR TITLE-ABS-KEY ( parasit* ) OR TITLE-ABS-KEY ( pathogen* ) OR TITLE-ABS-KEY ( zoono* ) OR TITLE-ABS-KEY ( microb* ) OR TITLE-ABS-KEY ( vir* ) OR TITLE-ABS-KEY ( protozo* ) OR TITLE-ABS-KEY ( mico* ) OR TITLE-ABS-KEY ( fungi* ) OR TITLE-ABS-KEY ( nematod* ) OR TITLE-ABS-KEY ( helmint* ) OR TITLE-ABS-KEY ( antibiotic* ) OR TITLE-ABS-KEY ( antimicrobial* ) OR TITLE-ABS-KEY ( amr ) OR TITLE-ABS-KEY ( antiparasit* ) OR TITLE-ABS-KEY ( drug* ) AND TITLE-ABS-KEY ( extensive* ) OR TITLE-ABS-KEY ( pastur* ) OR TITLE-ABS-KEY ( graz* ) ) )

( ( TITLE-ABS-KEY ( conventional* ) AND TITLE-ABS-KEY ( cow ) OR TITLE-ABS-KEY ( cattle ) AND TITLE-ABS-KEY ( milk ) OR TITLE-ABS-KEY ( dairy ) OR TITLE-ABS-KEY ( cheese ) AND TITLE-ABS-KEY ( infect* ) OR TITLE-ABS-KEY ( parasit* ) OR TITLE-ABS-KEY ( pathogen* ) OR TITLE-ABS-KEY ( zoono* ) OR TITLE-ABS-KEY ( microb* ) OR TITLE-ABS-KEY ( vir* ) OR TITLE-ABS-KEY ( protozo* ) OR TITLE-ABS-KEY ( mico* ) OR TITLE-ABS-KEY ( fungi* ) OR TITLE-ABS-KEY ( nematod* ) OR TITLE-ABS-KEY ( helmint* ) OR TITLE-ABS-KEY ( antibiotic* ) OR TITLE-ABS-KEY ( antimicrobial* ) OR TITLE-ABS-KEY ( amr ) OR TITLE-ABS-KEY ( antiparasit* ) OR TITLE-ABS-KEY ( drug* ) AND TITLE-ABS-KEY ( organic* ) ) )

Outputs generated by the two query strings were subjected to the same eligibility criteria and the same PRISMA procedure. The inclusion criteria were: (1) a publication must be an original study, in which conventional management was compared to organic management (CvsO), or intensive management was compared to extensive management (IvsE), of dairy cattle farms; (2) results must refer to: the presence, prevalence, or incidence of an infectious agent or infectious disease; the use of antimicrobials; the presence of drug-resistant (AMR) or drug-susceptible (AMS) microbes or parasites (where “drug” is intended as an antimicrobial product).

The exclusion criteria were as follows: publications classified as reviews, or other types of secondary studies such as book chapters, conference publications, notes and incomplete text articles. The full texts of all potentially relevant studies were downloaded in their entirety. A further round of screening was applied through careful reading of the abstracts (carried out by three co-authors independently, who subsequently discussed the doubtful cases), in order to collect and identify all those studies that met the eligibility criteria, gradually excluding those identified as “not pertinent for some classified reasons” ** as described in the PRISMA flow chart (Fig. 1).

**Figure 1.**
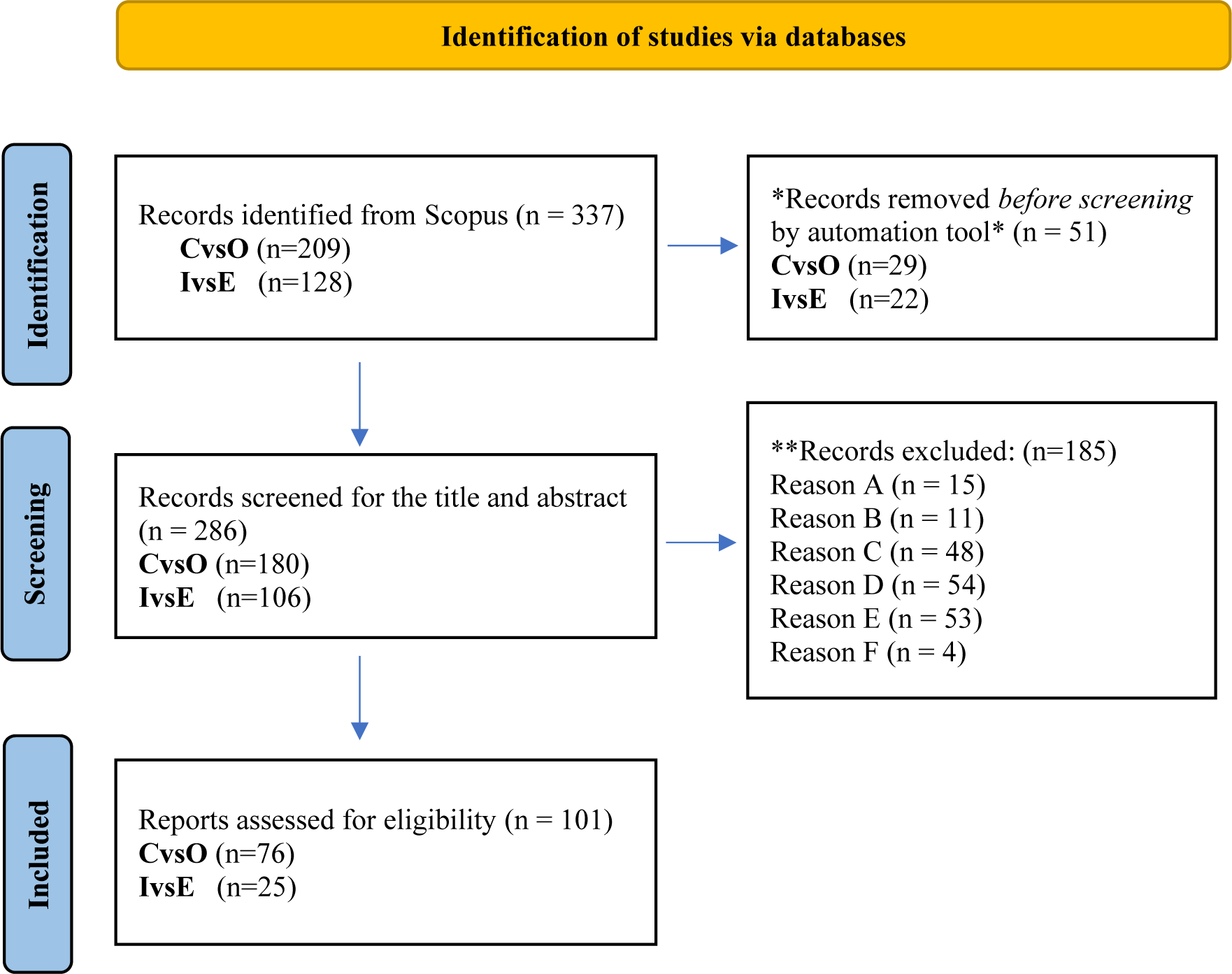
PRISMA flow chart presenting the results of the searching strategy and the exclusion process.* An automatic filter was applied to automatically exclude publications classified as “note”, “conference publications”, “book chapter”, “review”. ** A manual operation to identify publications defined “not pertinent” for the follow reasons (A, B, C, D, E, F); A: items focused on the issue of “consumer perception” or “farming policy”; B: items that did not refer to “dairy cow” (e.g., those focused on dairy buffalo or other dairy animals); C: items that did not perform, in the same study, a direct comparison CvsO or IvsE; D: the terms “extensive “intensive” “organic”, “conventional” did not refer to the farming management system; E: items that did not compare CvsO or IvsE according to eligibility criteria n° 2; F: publications classified as “note”, “conference publications”, “book chapter”, “review”.

## 3. RESULTS AND DISCUSSION

### 3.1. Overall results

A total of 336 items were retrieved from the Scopus platform. According to pre-defined eligibility criteria, as reported in the PRISMA flow-chart (Fig. 1), we finally selected 101 publications, 77 dealing with the comparison CvsO, and 25 with IvsE (one publication addressed both comparisons). We emphasize that part of these publications presented two or more studies, focused on different ID/IA or on both ID/IA and AMR. Therefore, the total number of studies (132) was higher than the number of publications (101). These 132 studies are reported in Tables 1-3, indicated as study numbers Sn1-Sn132, with the corresponding references. Also in the main text, the quoted studies will be referred to by indicating the Sn code, as reported in the Tables. The countries where the studies had been conducted and year of publication are reported in supplementary material (Tab S1 – S3). In total, 102 studies focused on f ID/IA and 30 on AMR. A higher number of studies was focused on CvsO (73 on ID/IA; 27 AMR), compared with IvsE (29 on ID/IA; 3 AMR). The countries where the studies had been conducted are also presented in figures 2 and 3 present. Comparative studies on IvsE were mainly conducted in Mediterranean and sub-Saharan countries, Far East Asia, and South America, while studies on CvsO were mainly conducted in North and South America, North and Central Europe, and Mediterranean countries. Studies on AMR/AMS came mainly from CvsO comparisons, mostly from North America and Europe. The two major ID groups that had been investigated in both CvsO and IvsE are: 1) intramammary infections (IMIs), with 31 studies (3 IvsE and 28 CvsO), mainly from North America and Europe and 2) gastrointestinal parasitic infections (GPIs), with 20 studies (4 IvsE and 16 CvsO) mainly from European countries. A full report and discussion on IMIs and GPIs can be found below in the main text. For the remaining IDs/IAs, for which the number of studies is limited, a complete list is reported below, with results and comments summarized in the supplementary material (Tab. S4). In general, the different dairy FMS (C, O, I, E) do not always appear to be associated with the incidence or prevalence of a given infection or disease. Furthermore, most of the studies did not clearly define the criteria for the attribution of a farm to the O or C group, even if several of the studies referred to the national transposition of the FAO guidelines on organic farms ^13^. Similarly, different studies classified I and E farms based on different criteria. Furthermore, a number of studies evaluated the risk factors associated with the prevalence/incidence of IDs, IAs or AMR, with less attention to the relevance of these risk factors in relation to the type of farming management systems (FMS) in terms of C, O, I, E. However, since the goal of this study was to contribute to the public debate on animal welfare and ecological sustainability issues associated with the different FMS, we will only report briefly on the specific risk factors (RFs).

**Fig. 2.**
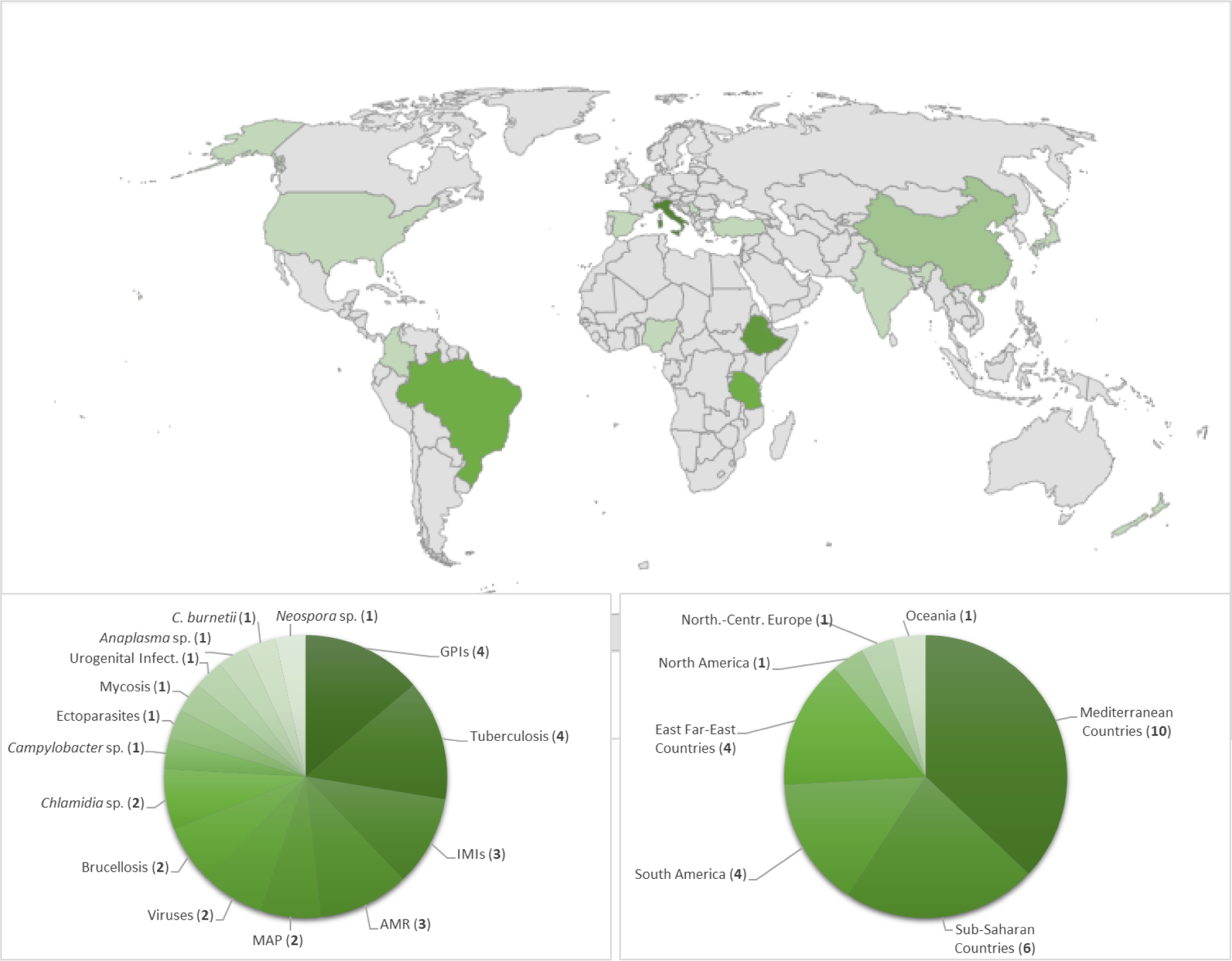
Countries where IvsE investigations have been conducted and relative prevalence of the studies on the different ID/IA or on AMR.

**Fig. 3.**
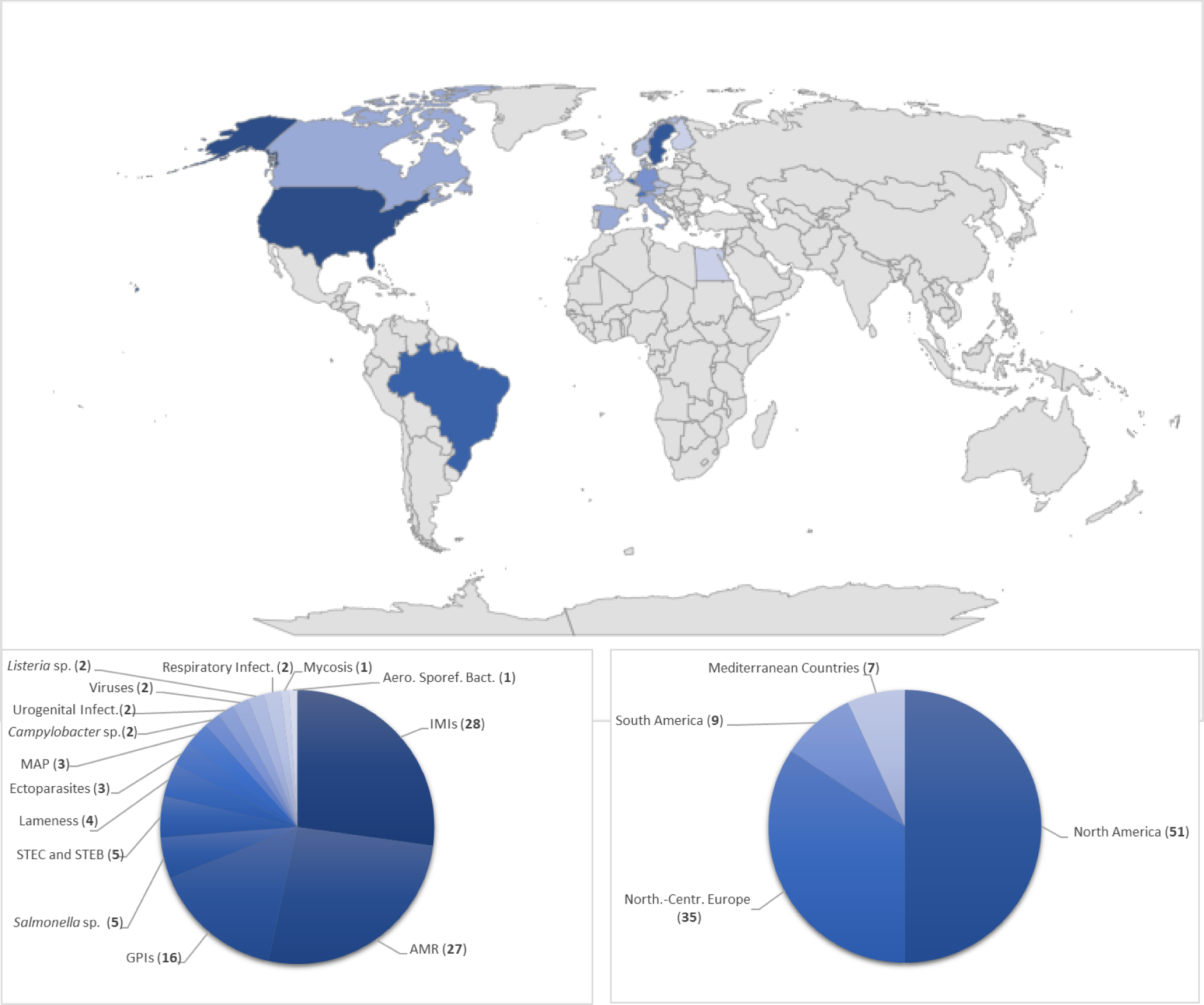
Countries where CvsO investigations have been conducted and relative prevalence of the studies on the different ID/IA or on AMR.

**Tab. 1.** Comparative studies on the presence of infectious diseases or infectious agents (ID/IA), in intensive versus extensive management (IvsE, on the left) and conventional versus organic management (CvsO, right), in which the investigated ID/IA coincide.

| ID/IA | IvsE |  | CvsO |  |
| --- | --- | --- | --- | --- |
|  | Sn | ref | Sn | ref |
| GPIs** | **Sn3-Sn6** | [57-60] | **Sn21-Sn36** | [14,18,25,40,45-56] |
| IMIs* | **Sn5-Sn7** | [42-44] | **Sn32-Sn59** | [14-41] |
| *Campylobacter* | **Sn8** | [72] | **Sn60, Sn61** | [70,71] |
| Ectoparasite | **Sn9** | [73] | **Sn62-Sn64** | [49,24,25] |
| MAP*** | **Sn10;Sn11** | [64,65] | **Sn65-Sn67** | [61-63] |
| Mycosis | **Sn12** | [74] | **Sn68** | [74] |
| Urogenital Inf. | **Sn13** | [69] | **Sn69, Sn70** | [29,34] |
| Viruses | **Sn14-Sn15** | [64,68] | **Sn71, Sn72** | [66,67] |

\*IMIs, Intra Mammary Infections; \*\*GPIs, Gastrointestinal Parasitic Infectious; \*\*\*MAP, *Mycobacterium avium* *subspecies paratuberculosis*, **Sn, study number**; ID/IA, infectious disease/infectious agent; ref, reference

**Tab. 2.**
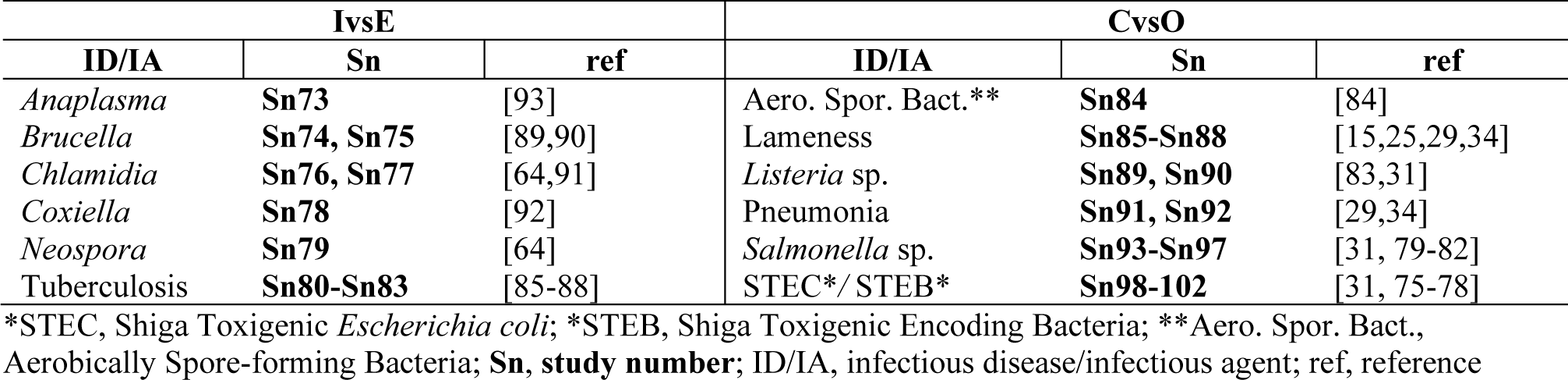
Comparative studies on the presence of infectious diseases or infectious agents (ID/IA), in intensive versus extensive management (IvsE, on the left) and conventional versus organic management (CvsO, right), in which the investigated ID/IA do not coincide.

**Tab. 3.** Comparative studies of antimicrobial resistance/antimicrobial susceptibility (AMR/AMS) in infectious diseases or infectious agents (ID/IA) in intensive versus extensive management (IvsE, on the left) and conventional versus organic management (CvsO, right)

| IvsE |  |  | CvsO |  |  |
| --- | --- | --- | --- | --- | --- |
| ID/IA | Sn | ref | ID/IA | Sn | ref. |
| *E. coli* | **Sn103** | [94] | *E. coli* | **Sn106-Sn114** | [19,76;96-103] |
| *Pasteurellaceae* | **Sn104** | [94] | *Campylobacter* | **Sn115, Sn116** | [113,71] |
| MCGT** | **Sn105** | [95] | IMIs*** | **Sn117-Sn125** | [30,32,39,41,104-108] |
|  |  |  | *Listeria* spp. | **Sn126** | [83] |
|  |  |  | MCGT** | **Sn127-Sn129** | [109-111] |
|  |  |  | *Salmonella* spp. | **Sn130, Sn131** | [19,112] |
|  |  |  | STEC* | **Sn132** | [98] |

\*STEC, Shiga Toxigenic *Escherichia coli*; \*\*MCGT, Microbial Community Gene Transfer; \*\*\*IMIs, Intra Mammary Infections; **Sn, study number**; ID/IA, infectious disease/infectious agent; ref, reference

### 3.2. Intramammary infections

Thirty-one studies focused on IMIs were retrieved. Twenty-eight of these dealt with CvsO (Sn32-Sn59) ^14–41^; three with IvsE (Sn5-Sn7) ^42–44^.

Eight out of the 28 studies on CvsO didn’t report any significative difference between C and O (Sn32; Sn35; Sn40; Sn44; Sn46; Sn49; Sn57; Sn58), six studies reported a higher prevalence/incidence of IMI in C (Sn38; Sn41; Sn42; Sn47; Sn52; Sn59); one study reported a higher prevalence/incidence of IMI in O (Sn56). Among the remaining 13 studies, six were focused on the risk factors associated with IMI (see below); seven highlighted apparently contrasting results. Sn55 recorded higher somatic cell counts (SCCs) in organic farms at day 31 DIM (day in milking), which were however similar to that of conventional herds at 102 DIM; in addition, higher prevalence of non-agalactiae streptococci was recorded at both 31 DIM and102 DIM in O compared to C, but the prevalence of coagulase-negative staphylococci was lower in O at 102 DIM. These fluctuations in cellular and microbial counts were not associated with clinical or sub-clinical mastitis. Similarly, in Sn41, higher counts of mesophilic and coliform bacteria were found in milk samples from O, but with no significant correlations with IMIs. A Norwegian study (Sn50) highlighted a higher proportion of dried off quarters in O vs. C, but did not find any difference in the number of quarters positive for mastitis bacteria, even if lower SCC in O cows was noted. Another study conducted in Sweden (Sn33) reported higher prevalence of IMI in C, even though bulk milk SCCs was higher in O. A German study (Sn51) didn’t report any difference in bulk milk SCCs, but reported a higher portion of organic vs. conventional individual cows with SCC > 150,000 at both 3 months before and 3 months after the dry period. A study conducted in the USA (Sn48) monitored several farms for three years during the transition from C to O, highlighting a higher incidence of mastitis in O at parturition, but no differences at dry-off. Finally, Sn39 reported that the incidence rate of clinical mastitis was higher in C compared to O (23.7 vs. 13.2 cases per 100 cows per year), however, bulk tank SCC tended to be lower in C.

Among the three retrieved studies on IvsE, in one the prevalence of mycotic mastitis in China was higher under extensive management (Sn6). One study, focused on *S. aureus* biofilm-producer strains in different regions of Serbia, didn’t find any differences under intensive or semi-extensive dairy farms (Sn5). Finally, a study conducted in the USA (Sn7) focused on the risk factors associated to grazing management.

Part of the studies retrieved with the query string for CvsO (namely: Sn34, Sn44, Sn47, Sn53, Sn54) were focused on the risk factors associated with IMI, and only to a lesser extent on the effects of the type of management, C or O, on the prevalence or incidence of these infections. Highlighted risk factors include: lactation number, farming part time, poor cleanliness of udders, size of the herd, use of mineral feed supplements, irregular milking intervals, milk urea concentrations, water temperature for washing the milking system, bedding area, timing of antibiotic treatments in relation to dry period (C only), hygiene, extent in the use of external resources, number of people responsible for mastitis treatment, age of the premises, percentage of cows with three or fewer quarters, use of fore stripping, proactive detection of mastitis during postpartum (and thus treatment), and stall barn housing. In addition, environmental temperature and duration of the infection in positive cows were associated with the incidence of mycotic mastitis (Sn6). Evidence for the role of genetic susceptibility to IMI was also reported in Sn36. A reduction in the risk of mastitis was associated to regular access to pasture, automatic milking shut-off and access to feed immediately after milking (Sn34). None of these factors appear to be specifically associated with C, O, I, E. However, they represent important leverage points for management towards healthier animals in terms of rational grazing (e.g., through rotational grazing rather than confined grazing) that are not specifically linked to intensive or extensive FM (Sn7).

### 3.3. Gastrointestinal parasitic infections

Twenty studies focused on GIPs were retrieved. Sixteen of these dealt with CvsO (Sn16-Sn31) ^14,45–47,18,48,49,25,50–54,40,55,56^, while the remaining four dealt with IvsE (Sn1-Sn4) ^57–60^.

Three out of 16 studies comparing CvsO (Sn21, Sn24, Sn25) didn’t report any significative difference, four studies reported a higher GPIs in conventional farming (Sn16, Sn18, Sn19, Sn30), five studies reported a higher GPIs in organic farming (Sn22, Sn23, Sn27, Sn29, Sn31), and the remaining four studies (Sn17, Sn20, Sn26, Sn28) were focused on risk factors associated to GPIs, not strictly related with the type of management. Studies on *Cryptosporidium* spp. infection provided apparently contradictory results. According to Sn24, there was no difference in the prevalence in either calves or cows; however, another study (Sn28) found higher levels of parasite shedding in organic farms, but a variety of factors not strictly related with the type of management might be associated with parasite shedding. Studies focused on fascioliasis (Sn21; Sn27), reported no significant difference between the two management types, while another (Sn18) detected significantly lower prevalence in O farms, probably due to continuous exposure to the parasite, leadings to better resilience (Sn16). Studies considering *Ostertagia ostertagi* highlighted contrasting results: two studies (Sn25; Sn29) found no correlation between infection and FMS, while another (Sn27) reported the opposite. Studies on the lungworm *Dictyocaulus viviparus* also reported contrasting results: Sn30 detected *D. viviparus* only on C, while Sn27 reported a prevalence of 18% in O and 9% in C herds. Interestingly, infected conventional herds were located near infected organic herds. Two studies (Sn22; Sn23), conducted in North and South America, reported a significatively higher prevalence of strongyle-type fecal eggs on O farms, while a German study (Sn19), highlighted a significantly higher prevalence C farms and pointed out the risk factors associated to seasonality. Another German study (Sn20) considered the issue of bovine genetics as a risk factor for parasitic infections, not strictly related to different FMS, even if particular genetic traits could be preferred in relation to the type of farming. Moreover, as highlighted in a Danish study (Sn17), estimations GPIs can also vary in relation with the type diagnostic method used.

Two out of four studies comparing IvsE reported higher GPIs on extensive farms (Sn2, Sn4), one reported higher GPIs in intensive farming, but also considered other risk factors (Sn1), while the other was focused only on risk factors (Sn3). Sn2 (conducted in Spain) reported a higher prevalence of *Ostertagia ostertagi* in EvsI, but the different environmental contexts could explain this result. Similar findings came from Sn4 (conducted in Italy) that compared two intensive farms (located in the Po River Valley) with an extensive farm (in an Alpine Mountain region), but again the differences in the sampling areas could represent an important confounding factor. Sn1, conducted in Ethiopia, highlighting a lower prevalence for cryptosporidiosis in IvsE; however, the authors note that the infection was significantly associated not only with the FMS but also with same others RFs like farm location, herd size, source of drinking water, weaning age, presence of bedding, pen cleanness and cleanness of hindquarters. Sn3, conducted in Tanzania, showed that prevalences of nematodes and flukes vary widely with geographic location and grazing management, further highlighting that several RFs not specifically related to the type of farming, (e.g. communal grazing and watering management practices) play a role in the circulation of parasitic worms.

### 3.4. Miscellaneous of retrieved IDs/IAs with limited comparative studies

This paragraph presents a complete list of the IDs/IAs for which the number of comparative studies is limited. Results and comments are summarized in the supplementary material (Tab S4).

IDs/IAs present on both CvsO and IvsE: five studies on MAP (Sn65-Sn67 CvsO ^61–63^; Sn10-Sn11 IvsE ^64,65^), four studies on viral infections (Sn71-Sn72 CvsO ^66,67^, and Sn14-Sn15 IvsE ^64,68^), three studies on urogenital infections (Sn69-Sn70 CvsO ^29,34^, and Sn13 IvsE ^69^), three studies on *Campylobacter* sp., (Sn60-Sn61 CvsO ^70,71^, and Sn8 IvsE ^72^), four studies on ectoparasites (Sn62-Sn64 CvsO ^49,24,25^, and Sn9 IvsE ^73^), two studies on zoonotic dermatophyte (Sn12 CvsO ^74^, and Sn68 IvsE ^74^).

IDs/IAs present only on CvsO: Shiga Toxigenic *E. coli* (STEC) and Shiga Toxigenic Bacteria (STEB) (Sn98-Sn102) ^31,75–78^, Lameness (Sn85-Sn88) ^15,25,29,34^, Salmonella sp. (Sn93-Sn97) ^31,79–82^, pneumonia (Sn91-Sn92) ^29,34^, Listeria sp. (Sn89-Sn90) ^83,31^ and aerobic spore-forming bacteria (Sn84) ^84^.

IDs/IAs present only on the IvsE: tuberculosis (Sn80-Sn83) ^85–88^, Brucellosis (Sn74-Sn75) ^89,90^, *Chlamidia* sp. (Sn76-Sn77) ^64,91^, *Coxiella burnetii* (Sn78) ^92^, *Neospora* sp. (Sn79) ^64^, and *Anaplasma marginale* (Sn73) ^93^.

### 3.5. Antibiotic resistance/susceptibility (AMR/AMS)

Among the 30 studies retrieved, 27 compared CvsO (Sn106-Sn132) and three IvsE (Sn103-Sn105). In general, a higher circulation of AMR genes was found on conventional and intensive FMS.

AMR/AMS in CvsO. Ten studies on AMR/AMS strains of *Escherichia coli* (Sn106-Sn114, Sn132) ^94,19,95–99,76,100,101^ were retrieved. Six were conducted in the USA. Sn113 didn’t find any significant difference across FMS for the abundance of *E. coli* O157 virulence marker genes, antimicrobial susceptibility profiles, and genotypes. On the contrary, Sn107 found significantly more abundant resistance genes in animals bred on conventional farms, but no significant differences in carcasses or beef trimmings. *E. coli* strains isolated from fecal samples showed a significantly higher resistance to seven out of 17 antimicrobial molecules on conventional farms (Sn114). Furthermore, AMR was strongly influenced by animal age, geographical region (dairy-intense flat land or more extensive foothill pasture) and whether cattle were raised for dairy or beef (Sn111). Sn112 assessed the association between age of cattle, and AMR/AMS in *E. coli* phylogroups isolated from both FMS. Here the authors used a hierarchical log-linear modeling approach that accounts for additional confounding factors, resulting in more robust relationships; The study provided evidence of clonal resistance (ampicillin) and genetic hitchhiking (tetracycline), reporting a significant association between low multidrug resistance, organic herds and numerically dominant phylogroup B1 strains, suggesting that the genetic composition of the herds may influence the AMR/AMS. Additionally, authors estimated that it would take from three to 15 years to have a significant change in bacterial populations passing from conventional to organic farming. Another study (Sn132) found a significantly higher proportion of non-susceptible spectinomycin in Shiga Toxigenic *E. coli* (STEC) isolates from conventional farms. Resistance to sulphadimethoxine in calves (but not in adult milking cows) was significantly higher on conventional farms. Multidrug resistant (MDR) patterns were more commonly found in non-O157 STEC vs. O157 STEC and the percentage of MDR on the two farm types was similar. A Swedish study (Sn108) found little significant difference for resistance to single antimicrobials on conventional farms. Most conventional herds had a rather high proportion of isolates resistant to at least one antimicrobial, but MDR strains were rare. A study conducted in Czech Republic (Sn110), revealed a higher prevalence of *E. coli* isolates producing an extended-spectrum beta-lactamase (ESBL) on conventional farms compared to organic ones. A Swiss study (Sn106) didn’t find any significative difference in the AMR of *E. coli* strains isolated from young dairy calves, but did report that the ESBL-producing Enterobacteriaceae were more prevalent in conventional farms.

Nine studies on AMR/AMS on IMIs were retrieved (Sn117-Sn125) ^102,103,30,32,104–106,39,41^. Three out the four studied conducted in USA (Sn125, Sn121, Sn122), found that *S. aureus* isolates from milk samples from C were significantly more resistant for the majority of tested antimicrobial molecules (Sn121, Sn125). Sn122 found that AMR of IMIs-associated pathogens were more common in C, yet the prevalence of bacteria responsible for mastitis was higher on O (with the exception of coliforms). Finally, a high proportion of sulfadimethoxine-resistant isolates were observed in both FMS, and were higher on O. Another study from USA (Sn119) was focused on the conversion process from conventional to organic management over a 3-year period; coagulase-negative staphylococci (CNS) were the most prevalent bacteria responsible for mastitis. They were significantly less resistant to β-lactam antibiotics after herd transitioned to O. AMR significatively decreased for ampicillin, cephalothin, cloxacillin, and penicillin for CNS, but not for *S. aureus*. This suggests that cessation of antibiotic use, in combination with organic management, reduced AMR of mastitis bacteria, even if the prevalence of *S. aureus* did not change significantly as herds transitioned from conventional to organic FMS. Similar results were reported in a Belgium study (Sn124) which found that the three most frequently isolated pathogens (*Streptococcus uberis*, *Staphylococcus aureus* and *Streptococcus dysgalactiae*) were significantly more resistant to antimicrobials on C. On the contrary, a Norwegian study (Sn120), did not find a significant difference in penicillin resistance against coagulase-negative staphylococci isolated from sub-clinically infected quarters, while a Swiss study (Sn123), found that AMR of *Staphylococcus* spp. and *Streptococcus* spp. was not significantly different in CvsO, except for *S. uberis,* which tended to have more single gene resistance on OvsC ones. Finally, a Brazilian study (Sn117), on *Staphylococcus* strains sampled in Minas Frescal cheese found no significant prevalence related to FMS.

Three studies on AMR/AMS in microbial communities and gene transfer phenomena MCGT (Sn127-Sn129) ^107–109^. A Canadian study (Sn127) analyzed publicly available metagenomics data, showing that organic practices are generally associated with lower prevalence of AMR Genes (AMRGs). Sn129 affirmed that the abundance and diversity of ARGs in feces was significantly higher in conventional vs. organic herds. All manure storage and soil samples had a diversity (albeit low abundance) of AMRGs conferring resistance to several antibiotics. Antimicrobial use on farms significantly influenced specific groups of AMRGs in feces, but not in manure storages or soil samples. Similar results from a Spanish study (Sn128), based on the monitoring AMRGs and mobile genetic elements (MGEs) in different types of comparisons (amended vs. unamended, CvsO, slurry vs. fresh or aged manure), found that the spread of AMRGs-MGEs cannot be inferred directly from any of the individual comparisons (including CvsO).

Two studies on AMR/AMS in *Salmonella* strains (Sn130, Sn131) ^19,110^. (both conducted in the USA). Sn130 found significantly higher circulation of AMRGs on C, but differences in carcasses or beef trimmings were not significant. Sn131, using logistic proportional hazards models, found that isolates from C were significantly associated with higher MIC for only two out of nine antimicrobials (streptomycin and sulfamethoxazole). Moreover, *Salmonella* isolates resistant to five or more antimicrobial agents were found on both FMS. Two studies on AMR/AMS conducted on *Campylobacter* strains (Sn115, Sn116) ^111,71^ (both conducted in the USA). Sn115 found that resistance to one out of four antimicrobial molecules tested was more prevalent in C, while Sn116, did not find any difference.

Similar results came from the only recovered study focused on *Listeria monocytogenes* (Sn126) ^83^, conducted in USA.

AMR/AMS in IvsE comparisons. Three studies on IvsE (Sn103-Sn105) ^112,113^.

An Indian study (Sn105) ^113^, comparing intensive farms vs. farms were animal had access to grazing, found that fecal bacteria from intensive farms were characterized by higher prevalence of AMR, which was also affected by feeding practices and nutrient concentration. Two studies, conducted by the same research group in Belgium (Sn103-Sn104) ^112^, focused on AMR profiles of both *E. coli*, (retrieved from the rectum) and *Pasteurellaceae* bacteria (retrieved from the nasal cavity) found a strong relationship between antimicrobial treatment and resistance profiles of bacterial isolates, in particular on intensive farms.

## 4. CONCLUSIONS

This systematic review has not specifically been focused on the issues of zoonoses, EID of human relevance, and spillover phenomena. Rather, we developed search strings with the objective of a wide-spectrum retrieval of research publications, dealing with the general issue of the circulation of infections and AMR in dairy farms in relation with the type of management system. However, results are relevant to the scenarios associated with zoonotic spillover events, an issue that has attracted a great deal of attention following the COVID19 pandemics ^114^. Indeed, the risks associated with intensive and extensive breeding systems are still debated (e.g., ^114^), and the present systematic review would suggest that the circulation of infectious agents is not significantly influenced by the type of management system in dairy farms, whether it is conventional or organic, intensive or extensive. Indeed, only a few studies reported significant differences, but with contrasting results regarding the higher or lower circulation of different pathogens under the different types of management. As anticipated in the introduction, the overall interpretation of our results suggest that CvsO and IvsE comparisons can be regarded as partially equivalent in some respects, for example in relation to the consumers’ perception of issues such as animal welfare and sustainability (e.g., ^115^ ^116^ ^117^). Therefore, the main outcome of our study, in relation to the public debate on the “pros” and “cons” of the different types of animal management, would suggest that organic and extensive farming is not correlated with a decrease of circulation of infectious agents compared to conventional and intensive dairy FMS.

Despite consumer perception that organic and extensive production are the same thing, the present study used two separate strings for literature search, in order to separate the two comparisons, CvsO and IvsE. In-depth examination of the retrieved publications highlight that: 1) the definitions of intensive and extensive farming differ among studies, making the comparison of studies that are apparently similar difficult; 2) the definitions of conventional and organic farming are often not stated in the publications, even if part of the studies refer to the national transposition of the general FAO guidelines on organic farming (which may differ from country to country) ^13^. Furthermore, different studies evaluated other risk factors, not necessarily associated with a given FMS.

Comparison between CvsO showed greater prevalence of antibiotic resistances in conventional farms. This is consistent with the minimal of antibiotics expected on organic farms, and the absence of selective pressure favoring the emergence of resistant strains. However, AMR was in general uncommon in both farming systems. Moreover, many confounding factors suggest that different bacterial species may behave differently depending on a variety of management and environmental variables in different environmental contexts and in different countries. For instance, age distribution, time of the year at sampling, an early or extended period of cow-calf contact, diagnostic methodology, type of drugs used, environmental context, etc. are quite different across the sampled farms, affecting interpretation of the significance. Herd size is particularly important because conventional herds maybe larger than organic ones or *vice-versa* and a reliable comparison is virtually impossible. As for the comparison IvsE, AMR was more prevalent in intensive farming, but results are to be interpreted with caution, since only three studies were retrieved for this comparison.

As already emphasized, the reading of the publications listed in Tables 1-3 revealed that terms like C, O, I, E are often defined differently. Furthermore, the results of the statistical tests in some of the examined studies might have been flawed by confounding factors, thus reducing the power of statistical analysis, which may have detected false significant correlations or not detected significant ones. For this reason, future studies on the impact of the management system on the circulation of infectious agents and AMR should apply statistical approaches exploiting generalized linear models (GLM) or similar statistical tools, instead of tests which only consider the variable of interest. Since GLMs account for additional factors, they can correctly identify the effect of confounding factors to highlight if significant differences really exist.

## Acknowledgments

Work supported by IRCAF - Invernizzi Reference Centre on Agri-Food “Romeo ed Enrica Invernizzi”. Project Milk Quality: a multifaceted approach. Massimo sincerely thanks Indira Fassioni for her kind support.

## SUPPLEMENTARY MATERIAL

**Tab. 1S:**
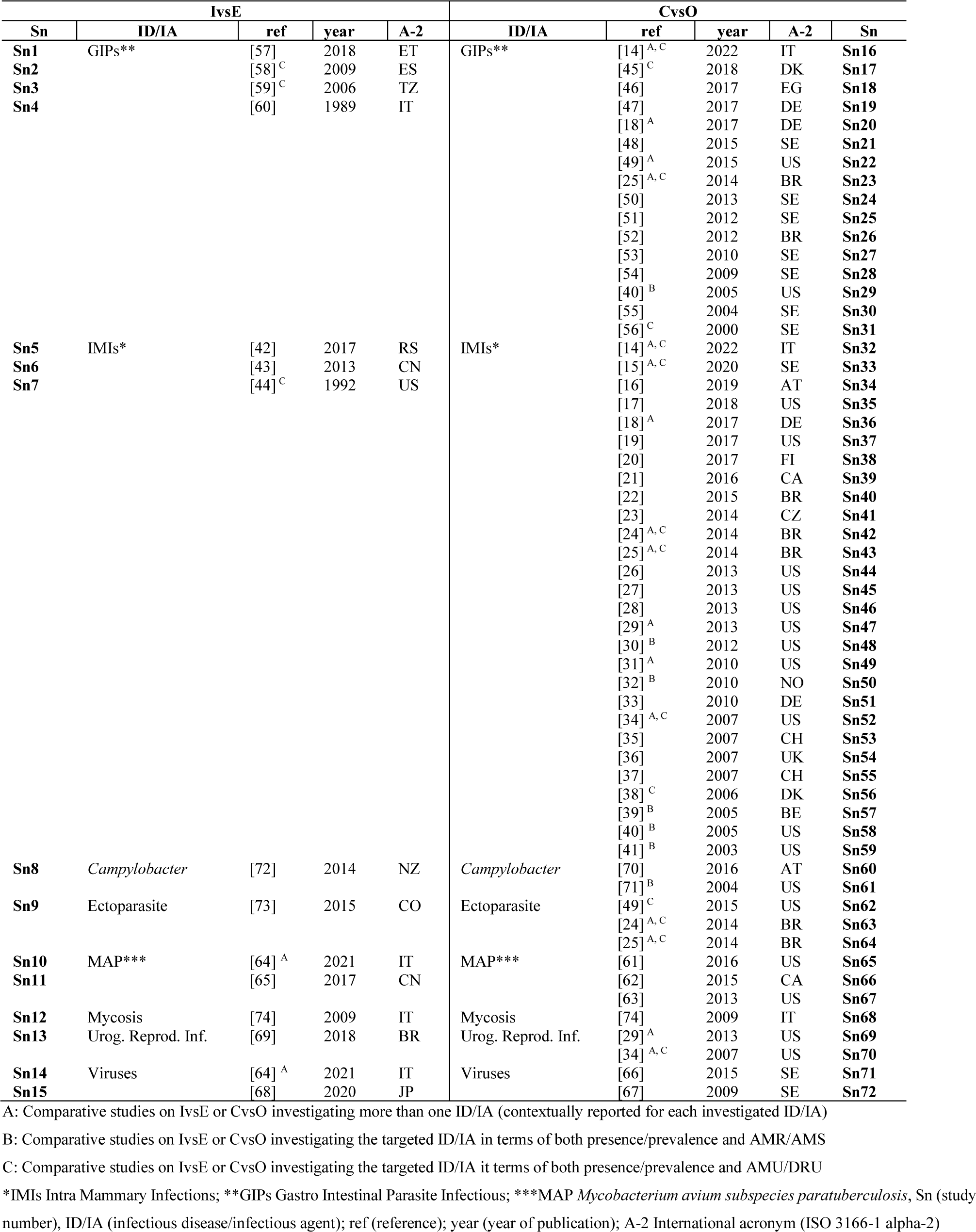
comparative studies on ID/IA presents in both comparations (CvsO, right) and (IvsE, on the left)

**Tab. 2S:** comparative studies on the presence of infectious diseases or infectious agents (ID/IA), in intensive versus extensive management (IvsE, on the left) and conventional versus organic management (CvsO, right), in which the investigated ID/IA do not coincide.

| IvsE |  |  |  |  | CvsO |  |  |  |  |
| --- | --- | --- | --- | --- | --- | --- | --- | --- | --- |
| Sn | ID/IA | ref | year | A-2 | ID/IA | ref | year | A-2 | Sn |
| Sn73 | *Anaplasma* | [93] | 2021 | TR | Aero. Spor. Bact.** | [84] | 2008 | BE | Sn84 |
| Sn74 | *Brucella* | [89] | 2017 | ET | Lameness | [15]<sup>A, C</sup> | 2020 | SE | Sn85 |
| Sn75 |  | [90] | 2016 | BR |  | [25]<sup>A, C</sup> | 2014 | BR | Sn86 |
| Sn76 | *Chlamidia* | [64]<sup>A</sup> | 2021 | IT |  | [29]<sup>A</sup> | 2013 | US | Sn87 |
| Sn77 |  | [91] | 2021 | BR |  | [34]<sup>A, C</sup> | 2007 | US | Sn88 |
| Sn78 | *Coxiella* | [92] | 1984 | NG | *Listeria* sp. | [83]<sup>B</sup> | 2022 | ES | Sn89 |
| Sn79 | *Neospora* | [64]<sup>A</sup> | 2021 | IT |  | [31]<sup>A</sup> | 2010 | US | Sn90 |
| Sn80 | Tuberculosis | [85] | 2012 | TZ | Pneumonia | [29]<sup>A</sup> | 2013 | US | Sn91 |
| Sn81 |  | [86] | 2009 | TZ |  | [34]<sup>A, C</sup> | 2007 | US | Sn92 |
| Sn82 |  | [87] | 2009 | ET | *Salmonella* sp. | [31]<sup>A</sup> | 2010 | US | Sn93 |
| Sn83 |  | [88] | 2003 | ET |  | [79] | 2005 | US | Sn94 |
|  |  |  |  |  |  | [80] | 2005 | US | Sn95 |
|  |  |  |  |  |  | [81] | 2005 | US | Sn96 |
|  |  |  |  |  |  | [82] | 2004 | US | Sn97 |
|  |  |  |  |  | STEC*/ STEB* | [31]<sup>A</sup> | 2010 | US | Sn98 |
|  |  |  |  |  |  | [75] | 2009 | US | Sn99 |
|  |  |  |  |  |  | [76]<sup>B</sup> | 2006 | US | Sn100 |
|  |  |  |  |  |  | [77] | 2006 | US | Sn101 |
|  |  |  |  |  |  | [78] | 2005 | CH | Sn102 |

A: Comparative studies on IvsE or CvsO investigating more than one ID/IA (contextually reported for each investigated ID/IA) B: Comparative studies on IvsE or CvsO investigating the targeted ID/IA in terms of both presence/prevalence and AMR/AMS C: Comparative studies on IvsE or CvsO investigating the targeted ID/IA it terms of both presence/prevalence and AMU/DRU \*STEC (Shiga Toxigenic *Escherichia coli*), \*STEB (Shiga Toxigenic Encoding Bacteria); \*\*Aero. Spor. Bact. (Aerobically Spore-forming Bacteria), Sn (study number); ID/IA (infectious disease/infectious agent); ref. (reference); year (year of publication), A-2 International acronym (ISO 3166-1 alpha-2)

**Tab. 3S:** comparative studies of antimicrobial resistance/antimicrobial susceptibility (AMR/AMS) in infectious diseases or infectious agents (ID/IA) in intensive versus extensive management (IvsE, on the left) and conventional versus organic management (CvsO, right)

| IvsE |  |  |  |  | CvsO |  |  |  |  |
| --- | --- | --- | --- | --- | --- | --- | --- | --- | --- |
| Sn | ID/IA | ref | year | A-2 | ID/IA | ref | year | A-2 | Sn |
| Sn103 | *E. coli* | [94]<sup>A</sup> | 2016 | BE | *E. coli* | [96] | 2022 | CH | Sn106 |
| Sn104 | *Pasteurellaceae* | [94]<sup>A</sup> | 2016 | BE |  | [19]<sup>A</sup> | 2020 | US | Sn107 |
| Sn105 | MCGT** | [95] | 2023 | IN |  | [97]<sup>C</sup> | 2020 | SE | Sn108 |
|  |  |  |  |  |  | [98] | 2015 | PL | Sn109 |
|  |  |  |  |  |  | [99] | 2011 | CZ | Sn110 |
|  |  |  |  |  |  | [100] | 2010 | US | Sn111 |
|  |  |  |  |  |  | [101] | 2007 | US | Sn112 |
|  |  |  |  |  |  | [76]<sup>B</sup> | 2006 | US | Sn113 |
|  |  |  |  |  |  | [102] | 2005 | US | Sn114 |
|  |  |  |  |  | *Campylobacter* | [113]<sup>B</sup> | 2006 | US | Sn115 |
|  |  |  |  |  |  | [71]<sup>B</sup> | 2004 | US | Sn116 |
|  |  |  |  |  | IMIs*** | [104] | 2021 | BR | Sn117 |
|  |  |  |  |  |  | [105] | 2018 | DE | Sn118 |
|  |  |  |  |  |  | [30]<sup>B</sup> | 2012 | US | Sn119 |
|  |  |  |  |  |  | [32]<sup>B</sup> | 2010 | NO | Sn120 |
|  |  |  |  |  |  | [106] | 2008 | US | Sn121 |
|  |  |  |  |  |  | [107]<sup>C</sup> | 2007 | US | Sn122 |
|  |  |  |  |  |  | [108] | 2006 | CH | Sn123 |
|  |  |  |  |  |  | [39]<sup>B</sup> | 2005 | BE | Sn124 |
|  |  |  |  |  |  | [41]<sup>B</sup> | 2003 | US | Sn125 |
|  |  |  |  |  | *Listeria* spp. | [83]<sup>B</sup> | 2022 | ES | Sn126 |
|  |  |  |  |  | MCGT** | [109] | 2023 | CA | Sn127 |
|  |  |  |  |  |  | [110] | 2021 | ES | Sn128 |
|  |  |  |  |  |  | [111] | 2020 | US | Sn129 |
|  |  |  |  |  | *Salmonella* spp. | [19]<sup>A</sup> | 2020 | US | Sn130 |
|  |  |  |  |  |  | [112] | 2006 | US | Sn131 |
|  |  |  |  |  | STEC* | [103] | 2007 | US | Sn132 |

A: Comparative studies on IvsE or CvsO investigating more than one ID/IA (contextually reported for each targeted ID/IA) B: Comparative studies on IvsE or CvsO investigating the targeted ID/IA it terms of both, presence/prevalence and AMR/AMS C: Comparative studies on IvsE or CvsO investigating the targeted ID/IA it terms of both, presence/prevalence and AMU/DRU \*STEC (Shiga Toxigenic *Escherichia coli*); \*\*MCGT (Microbial Community Gene Transfer); \*\*\*IMIs Intra Mammary Infections, Sn (study number), ID/IA (infectious disease/infectious agent); ref (reference); year (year of publication); A-2 International acronym (ISO 3166-1 alpha-2)

**Tab. 4S:**
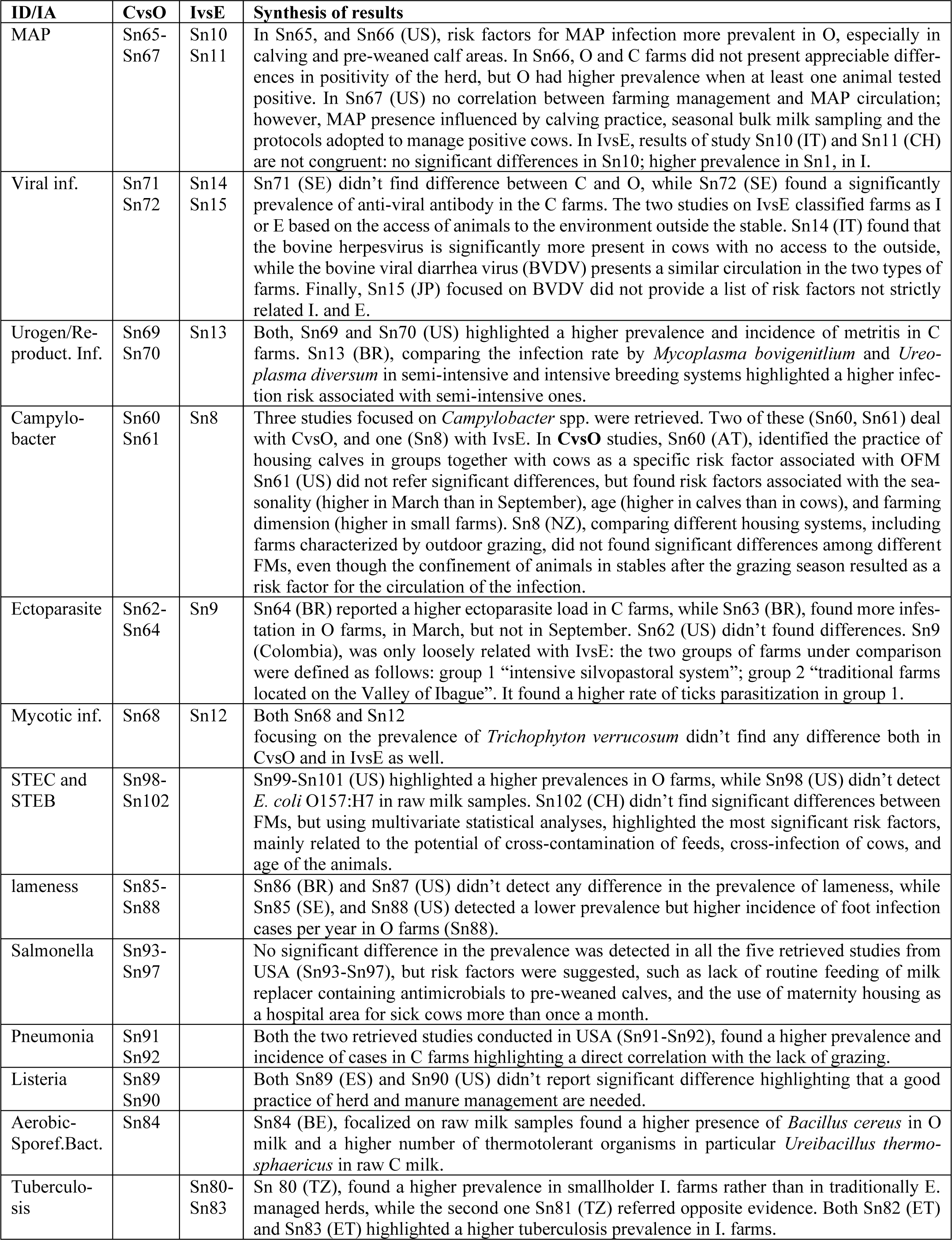
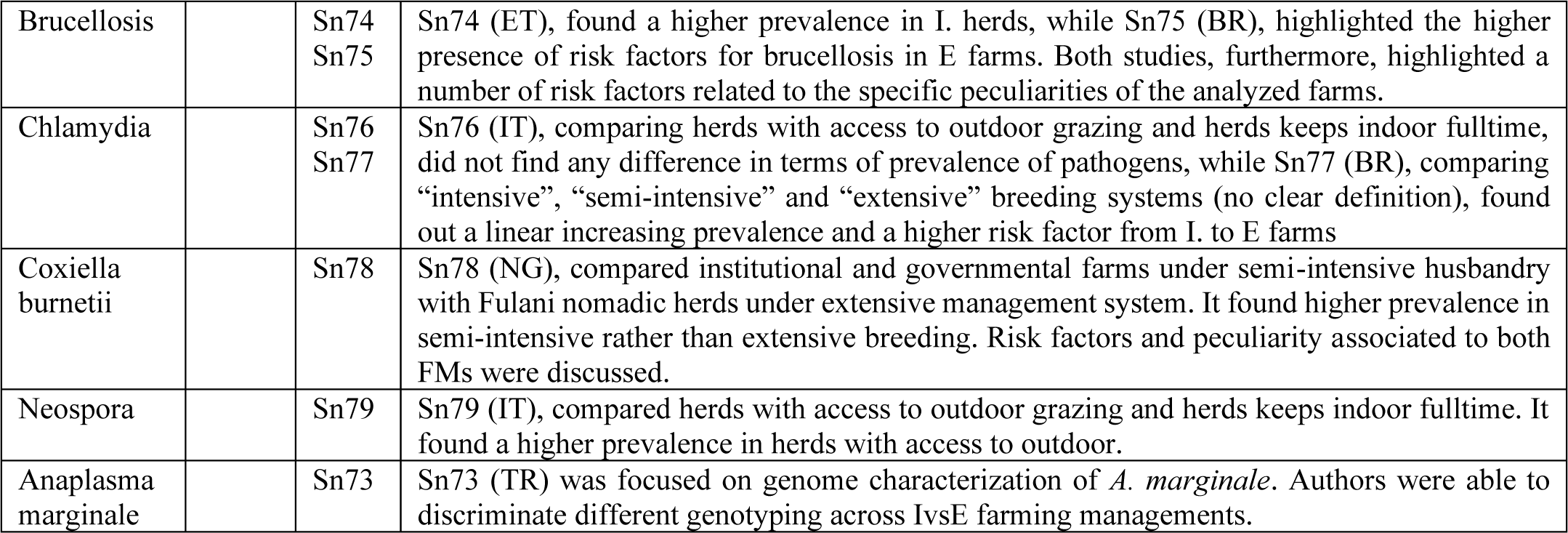
Results and comments for the IDs/IAs with limited number of comparative studies.

